# SOX4-SMARCA4 complex promotes glycolysis-dependent TNBC cell growth through transcriptional regulation of Hexokinase 2

**DOI:** 10.1101/2023.09.10.557071

**Authors:** Pooja Khanna, Rushabh Mehta, Gaurav A. Mehta, Vrushank Bhatt, Jessie Y. Guo, Michael L. Gatza

**Author notes:** Correspondence: Michael L. Gatza, PhD., Rutgers Cancer Institute of New Jersey, 195 Little Albany Street CINJ 4558, New Brunswick, NJ 08903.

## Abstract

Tumor cells rely on increased glycolytic capacity to promote cell growth and progression. While glycolysis is known to be upregulated in the majority of triple negative (TNBC) or basal-like subtype breast cancers, the mechanism remains unclear. Here, we used integrative genomic analyses to identify a subset of basal-like tumors characterized by increased expression of the oncogenic transcription factor SOX4 and its co-factor the SWI/SNF ATPase SMARCA4. These tumors are defined by unique gene expression programs that correspond with increased tumor proliferation and activation of key metabolic pathways, including glycolysis. Mechanistically, we demonstrate that the SOX4-SMARCA4 complex mediates glycolysis through direct transcriptional regulation of Hexokinase 2 (HK2) and that aberrant HK2 expression and altered glycolytic capacity are required to mediate SOX4-SMARCA4-dependent cell growth. Collectively, we have defined the SOX4-SMARCA4-HK2 signaling axis in basal-like breast tumors and established that this axis promotes metabolic reprogramming which is required to maintain tumor cell growth.

## INTRODUCTION

Triple negative breast cancer (TNBC) is an aggressive malignancy characterized by increased proliferation and metastatic capacity, limited therapeutic options and poor clinical outcome [1–4]. These tumors are largely synonymous with the basal-like molecular subtype and lack expression of identified therapeutic targets including estrogen receptor, progesterone receptor and human epidermal growth factor receptor. As such, limited therapeutic options are available for these patients beyond cytotoxic chemotherapy, radiation and surgery [1, 2, 5–7].

In addition to the genomic heterogeneity that exists between clinical and molecular subtypes of breast cancer, metabolic heterogeneity is evident among tumor types and subtypes and contributes to tumor development and progression [8, 9]. Rapidly dividing tumor cells undergo metabolic reprogramming to take advantage of available nutrients in order to synthesize the necessary macromolecules and metabolites required to support sustained cell proliferation, tumor growth and disease progression, including the emergence of therapeutic resistance [8–11]. Consistent with these general observations, a number of studies have demonstrated that TNBC or basal-like breast tumors are characterized by increased expression of glycolytic enzymes and increased dependency on glycolysis for cell viability [12–14]. Despite these observations, the underlying mechanisms that promote glycolysis in these tumors remain unclear.

The oncogenic transcription factor SOX4 (SRY-related-HMG box 4) is highly expressed, and can mediate tumor development and progression, in multiple forms of cancer, including basal-like breast tumors [15–20]. SOX4 mediates tumorigenic activity through aberrant activation of multiple oncogenic pathways, including PI3K, Wnt/β-catenin, TGFβ, and others, which collectively contribute to changes in proliferation, cell cycle, EMT, stemness and angiogenesis [15, 21–24]. We recently reported that SOX4 forms a complex with the SWI/SNF ATPase SMARCA4 (SS complex) which is required to mediate SOX4-dependent activation of TGFβ and PI3K signaling in basal-like breast cancer [24]. Similar to SOX4, SMARCA4 has been reported to correlate with poor prognosis and to contribute to tumorigenesis by modulating critical aspects of breast cancer biology including lipid metabolism, proliferation and chemotherapy resistance [25–29]. While the SS complex is highly expressed in basal-like breast tumors, the global impact of this complex on cellular signaling and tumorigenesis remains unknown.

The goal of this study was to define the SOX4-SMARCA4 (SS) complex gene expression program and to determine the impact of this signaling network on TNBC tumorigenesis. We determined that basal-like tumors with high SS complex expression are defined by unique gene expression programs associated with increased proliferation and glycolysis. SOX4 and SMARCA4 expression promotes metabolic reprogramming and mechanistic studies demonstrated the effect of the SOX4-SMARCA4 complex on cell growth and glycolysis. We determined that SOX4 regulates Hexokinase 2 (HK2) expression in a SMARCA4 dependent manner, and that HK2-dependent glycolysis is required to mediate SS complex-dependent cell growth. This study established the SOX4-SMARCA4 complex as an essential factor that promotes metabolic reprogramming and proliferation in basal-like tumors.

## MATERIALS AND METHODS

### Gene expression data

Gene expression data were acquired for human breast samples from The Cancer Genome Atlas (TCGA) (n=1,032) [30] and Molecular Taxonomy of Breast Cancer International Consortium (METABRIC) (n=1,992) [31] data portals and processed as previously described [24]. Cancer Cell Line Encyclopedia (CCLE) RNAseq and mass spectrometry data were acquired for basal-like breast cancer cell lines (https://portals.broadinstitute.org/ccle) [32].

### SOX4-SMARCA4 subgroup analyses

Gene expression data for basal-like tumors from TCGA (n=185) and METABRIC (n=331) were extracted based on PAM50 subtypes [30, 33]. A Pearson correlation was used to identify positively (r>0.2) or negatively (r<-0.2) correlated genes (p<0.01) in the TCGA dataset relative to both SOX4 and SMARCA4 expression then validated in the METABRIC dataset to identify a consensus gene set (Table S1). Consensus Cluster Plus [34] was used to classify basal-like tumors based on the consensus gene list (Table S2 and Table S3).

Two-class SAM analysis [35] was used to identify differentially expressed genes, methylation and protein expression in SS high and low basal-like clusters using data from TCGA (n=185). GSEA analysis [36] was used to identify pathways of interest using genes with consistent gene expression (q< 0.05) and methylation (q<0.01) patterns. SAM analyses of Reverse Phase Protein Array (RPPA) data (TCGA, n=179) from basal-like tumor samples were used to compare protein expression in SS complex high and low tumors, including a previously published protein expression signature [37]. TNBC subtype was assigned using the TNBCtype tool (https://cbc.app.vumc.org/tnbc) (Tables S2 and S3) and the relationship between TNBC subtype and SS cluster statistically assessed by a Fisher’s exact test [38, 39]. A Spearman rank correlation was used to examine the relationship between average SOX4 and SMARCA4 expression and proliferation or glycolysis gene expression signatures [40–42].

### SOX4 interactome analyses

Cytoscape (version 3.8.0) was used to visualize associations between SOX4 and SWI/SNF components from mass spectrometry analysis [24]; edge length was the inverse of the ratio of the spectral counts (IgG vs SOX4-V5).

### Breast cancer cell lines and gene manipulation

HCC1143, HCC1954, MDA-MB-468, HCC1395, HCC38, HCC70, BT20, MDA-MB-231 and MCF10A cell lines were purchased from ATCC and cultured according to ATCC guidelines. For overexpression studies, the following lentiviral vectors were used at an MOI of 4 (96h): pWPXL-SOX4 (Addgene 36984), HA-tagged SOX4 [43] or empty vector control (Addgene 12257). SMARCA4 was overexpressed by WT SMARCA4-sfGFP (Addgene 107056) MOI:3 (96h). For knockdown studies, shRNA against SOX4 (sh1: TRCN000018213, sh2: TRCN000018214) (Dharmacon), HK2 (RHS4531-EG3099, Dharmacon) or a scramble control (VSC11649) were used (MOI:3, 72-96h). Cells were plated at a density of 100,000 (MCF10A) or 300,000 (MDA-MB-468 or HCC1954) cells/10cm^2^ plate then transduced with 8μg/ml polybrene 24h after seeding. HK2 was overexpressed with the FLHKII-pGFPN3 vector (Addgene 21920) at 10µg/10cm^2^ dish for 96h using Lipofectamine 2000 (Thermo Fisher). To inhibit SMARCA4 expression, 300,000 cells were transfected with 50nm ON-TARGETplus Human SMARCA4 siRNA (Dharmacon L-010431-00-0005) or the corresponding siControl (D0012061305) by lipofectamine RNAiMAX (ThermoFisher) according to the manufacturer’s protocol. Changes in gene or protein expression were verified by qRT-PCR or Western blot.

### RNA extraction and quantitative real-time PCR

Total mRNA was isolated using the RNeasy Plus Mini Kit (Qiagen 74136) and cDNA was synthesized using the QuantiTech Reverse Transcription Kit (Qiagen 205311) following the manufacturer’s instructions. The qPCR was conducted using 12.5µl SYBR Select Master Mix (Applied Biosystems 4472919), 10µM of each primer (IDT), 10.3µl RNase free water and 2µl cDNA per reaction for a total volume of 25µl. The qPCR was performed using an Applied Biosystems QuantStudio 3 Real-Time PCR System (Version 1.4) with the following primers: human *SOX4* Forward: 5’-CTCTCCAGCCTGGGAACTATAA-3’, *SOX4* Reverse 3’- CGGAGGTGGGTAAAGAGAGAA-5’; human *Beta-Actin* Forward 5’- GCACCACACCTTCTACAATG-3’, Reverse 3’-TGCTTGCTGATCCACATCTG5-5’; human *SMARCA4* Forward 5’-AGTGCTGCTGTTCTGCCAAAT-3’, Reverse 3’- CCGTCGTTGAAGGTTTTCAG-5’; and human *HK2* Forward 5’- CAGCTATTTGGGAGGCTGAG-3’, Reverse 3’-TAACTGGGCTTCCCTCTTCA-5’.

### Protein lysates and Western blot analyses

Cells were harvested in Triton Lysis Buffer (25mm HEPES, 100mm NaCl, 1mm EDTA, 10% glycerol, and 1% Triton X-100) with 1x protease and phosphatase inhibitor (Cell Signaling Technology, 5872S) and protein concentration determined by Pierce BCA Protein Assay Kit (23225). For Western blot analyses, 35-50µg were loaded on a 4-20% TGX Gradient Gel (BioRad) and run at 100V for 1.5 hours at room temperature. Gels were transferred onto a PVDF membrane overnight at 35V at 4LJC. Membranes were blocked with Advanblock blocking buffer (Advansta R-03726-E10) for 1 hour at room temperature and incubated at 4°C in primary antibody overnight. The signal was developed with the SuperSignal West Pico Plus Chemiluminescent Substrate (ThermoFisher 34580) and imaged on the ChemiDoc Touch Imaging System (BioRad). The antibodies were diluted in Advanblock buffer at the following concentrations: SOX4 1:500 (Boster Biological Technology PB9618), HK2 1:1000 (CST, 2867), SMARCA4 1:1000 (SantaCruz, SC-374197), Beta-Actin 1:10000 (CST, 4970S), HA 1:5000 (CST, C29F4 3724), PCNA 1:1000 (CST, 13110), cleaved PARP (CST, 5625) PARP 1:1000 (CST, 9542) and Histone H3 (CST 4499).

### Cell proliferation assays

Cell proliferation colorimetric-based assays were performed using 4,000 transduced or transfected cells and the CellTiter 96 Aqueous One Solution Cell Proliferation Assay (Promega, G3582) according to the manufacturer’s instructions for 96h. For cell growth assays, 32,500 MCF10A cells were transduced with empty vector, SOX4 and/or SMARCA4 lentiviruses and cell growth was assessed after 7 days by 0.05% crystal violet for 10 minutes at room temperature followed by two washes in distilled water. Plates were dried then imaged and quantified by ImageJ. For drug treatment assays, 3-bromopyruvic acid (MedChemExpress, HY-19992) was diluted in DMSO and cells were treated at the indicated concentrations. Prior to treatment with the given concentrations of 3-bromopyruvate, 32,500 MCF10A cells were transduced to express EV or SOX4 and SMARCA4 lentivirus (MOI:4), the cells were allowed to grow for 5 days and were then seeded at a density of 100,000 cells per well of a 24 well plate and allowed to recover for 24 hours. Cells were then treated with drug for 24 hours, stained and quantified as described above.

### Colony Formation Assay

For colony formation assays, cells were transduced using gene-specific shRNAs or transfected by gene-specific siRNAs or scrambled control as outlined. After 48 hours cells were transduced again. 24 hours after the second transduction, 4,000 cells were seeded in a 6 well plate, and grown for a total of 10 days with medium changes every 48 hours. For the SOX4 and SMARCA4 overexpression assays, cells were transduced as described above, medium was changed the following day, then cells were plated in a 24 well plate and quantified as above, for a total experimental time of 7 days. At either endpoint, cells were stained by crystal violet and imaged as described above. Images were quantified by intensity using ImageJ Fiji software for at least three independent experimental replicates and normalized to control.

### Metabolic and metabolite analyses

MCF10A cells were transduced with pWPXL-SOX4 or empty vector control (MOI:4, 96h). Cell medium was then aspirated, cells were washed three times in sterile PBS and medium lacking sodium pyruvate with dialyzed serum (DMEM ThermoFisher MT-10-017-CV and SH3007903) added for 24 hours. Two hours prior to sample harvest, the medium was replaced with fresh medium. To harvest lysates, cells were washed with sterile PBS once, ice cold 1ml 40:20:20 methanol:acetonitrile:water with 0.5% formic acid was added to the plate and cells were incubated on ice for 5 minutes. After incubation, 50µl 15% NH_4_HCO_3_ was added, cells were scraped and centrifuged at 15000g for 10 minutes at 4LJC. The supernatant was aspirated and the cell pellet stored at −80LJC. Metabolite levels were analyzed by liquid chromatography-mass spectrometry for metabolites as previously described [44].

For ^13^C tracing experiments, MDA-MB-468 cells were transduced with a scrambled control or shSOX4 lentivirus. 300,000 cells were seeded to a new 10cm^2^ plate for 72 hours. To label cells, ^13^C glucose (Cambridge Isotope Libraries 110187-42-3) was added to media lacking sodium pyruvate and glucose (Gibco 11966) for 3 minutes to measure glycolytic flux. The 6^th^ carbon was measured for glucose, G6P and F6P and the 3^rd^ carbon for all other glycolytic pathway components. For metabolite analyses by MS, measurements were normalized to cell counts. All measurements were normalized to the average of control cells.

### Glycolytic stress test

Seahorse glycolytic stress test was performed according to the manufacturer’s protocol. Briefly, MCF10A cells were seeded at a density of 100,000 cells on a 10cm^2^ plate. Twenty-four hours later, cells were transduced to overexpress either wild-type SOX4 or HA-tagged SOX4 (MOI:4) or transfected to inhibit SMARCA4 expression by siRNA. After 72 hours, 5,000 cells from each experimental condition were seeded in 100μl MCF10A medium per well in triplicate on a Seahorse 24-well plate. After 6 hours, an additional 1ml of medium was added to each well and cells were allowed to grow for 18 hours. To perform the glycolytic stress test, cells were incubated for 1 hour in Seahorse medium (RPMI without glucose or pyruvate). Extracellular Acidification Rate (ECAR) was measured using the XF24e Seahorse Biosciences Flux Bioanalyzer. Three cycles (3 minutes mixing, 2 minutes waiting and 3 minutes measurement) were conducted for each of the four stages of experiment: basal ECAR, glycolysis, glycolytic capacity and glycolytic reserve. Basal glycolytic rates were measured for the first 24 minutes. After 24 minutes, 10mm glucose was injected into each well and the shift in ECAR levels was measured for 24 minutes to determine the rate of glycolysis. 1μM of the ATP synthase inhibitor oligomycin was then injected to inhibit mitochondrial ATP production to measure maximum glycolytic capacity (24 minutes). Finally, 50mm of 2-deoxy glucose (2-DG) was injected into each well to inhibit glycolysis (24 minutes). Results were analyzed using the Wave software and a t-test used to establish statistical significance.

### *In vitro* glucose and glucose-6-phosphate assays

Intracellular glucose (Abcam AB65333) and glucose-6-phosphate (Abcam AB83426) levels were measured using colorimetric assays following SOX4, HK2 or SMARCA4 manipulation in MCF10A, MDA-MB-468 or HCC1954 cell lines. Cells were harvested by trypsinization at 72h for the knockdown experiments and 96h for the overexpression experiments. To measure metabolites, medium was aspirated, cells were washed in sterile 1x PBS to remove extracellular glucose or glucose-6-phosphate and centrifuged at 4,000 rpm for 1 minute at 4LJC. Cells were counted and 50,000 cells resuspended in the supplied glucose or glucose-6-phosphate buffer and seeded in triplicate in a 96 well plate. Cells were incubated with the respective reaction mix for each kit, which contained 2µl enzyme, 2µl probe and 46µl buffer per well in the dark for 30 minutes at 37°C and a Tecan Infinite F50 used to examine glucose (570nm) or glucose-6-phosphate (450nm) levels. Measurements were then calculated relative to a glucose or glucose-6-phosphate standard curve and normalized to control.

### ChIP PCR

Chromatin Immunoprecipitation (ChIP)-PCR analyses were conducted as described [24, 45]. MCF10A cells were transduced for 96 hours with the HA tagged SOX4 construct (MOI:4) as described above, then crosslinked with formaldehyde for 10 minutes at room temperature. The reaction was quenched with 2.5M glycine for 5 minutes at room temperature, then washed and resuspended in buffer containing 50mm HEPES-KOH, pH 7.5, 140mm NaCl, 1mm EDTA, 10% glycerol, 0.5% NP40, 0.25% Triton X-100, and phosphatase/protease inhibitor (Cell Signaling Technology, 5872S). Nuclei were resuspended in 10mm Tris-HCl, pH 8.0, 200mm NaCl, mm EDTA, 0.5mm EGTA and phosphatase/protease inhibitor, then lysed in 10mm Tris-HCl, pH 8.0, 100mm NaCl, 9mm EDTA, 0.5mm EGTA, 0.1% Na-Deoxycholate, 0.5% N-lauryolsarcosine and 1.1% Triton X-100. Lysates were pulled down with either anti-HA Magnetic Beads (VWR PI88836) or IgG (Invitrogen MA5-14453) then Protein G beads (Thermofisher 10004D). Lysates were washed 5 times in RIPA buffer and once with TE buffer containing 50mm NaCl. Complexes were eluted in 10mm Tris (pH 8.0) and assessed using the following primers and the qPCR protocol described above: human *HK2* promoter Forward 5’- TCAGAGGCAGAAGAACCACA-3’, Reverse 3’-GAGCTTGCAGTGAGCAGAGA-5’ and negative control human *PNOC* Forward 5’-GCTTGAGCTCCTTGGATGAC-3’ and Reverse 5’- CCTGTCCCTTACTGCAGA-3’. For analysis, the samples run in triplicate were averaged, the negative control PNOC values subtracted, these values were then subtracted from the IgG control and finally normalized to the empty vector control. To verify equal SMARCA4 pulldown in control or SOX4 siRNA treated HCC1954 cells, immunoprecipitation (IP) followed by Western blot analyses were performed using 5µg SMARCA4 or IgG antibodies as outlined above, lysates were washed 3 times and protein eluted with 2x SDS buffer and SMARCA4 and histone H3 assessed by Western blot.

### Analysis of gene expression and pathways in basal-like tumor samples

To examine the relationship between SOX4, SMARCA4, and HK2 expression in basal-like tumors, TCGA or METABRIC samples were stratified in into SOX4 high (top quartile) or low (all others) and a t-test used to assess significance. To examine glycolysis, glucose depletion or proliferation gene expression signatures [40–42], tumors were scored as previously reported and a t-test used to assess differences. To plot the data, genes were median centered and heatmaps were generated using JavaTreeView (Version 1.1.6r4) [46]. A Pearson correlation was used to demonstrate the relationship between the average of SOX4 and SMARCA4 expression and proliferation and glycolysis gene expression signatures in the TCGA and METABRIC datasets; data were visualized using the scatter3 function in MATLAB (version R2020a).

### Statistical analysis

For each experiment, at least three independent experiments were quantified as indicated and a t-test used to assess significance, except where noted.

## RESULTS

### Characterization of SOX4-SMARCA4 expressing basal-like breast tumors

We recently reported that the oncogenic transcription factor SOX4 forms a complex with the SWI/SNF ATPase SMARCA4, that both proteins are highly expressed in basal-like breast tumors and that this complex is required to regulate TGFβ and PI3K/Akt signaling [24]. Beyond the interaction with SMARCA4, our analyses of the SOX4 interactome using data from SOX4 immunoprecipitation followed by mass spectrometry (IP-MS) [24] identified strong protein-protein interactions between SOX4 and multiple SWI/SNF complex proteins (Figure 1A), with the strongest interaction between SOX4 and SMARCA4, leading us to hypothesize that this complex may modulate additional aspects of basal-like breast cancer biology.

**Figure 1.**
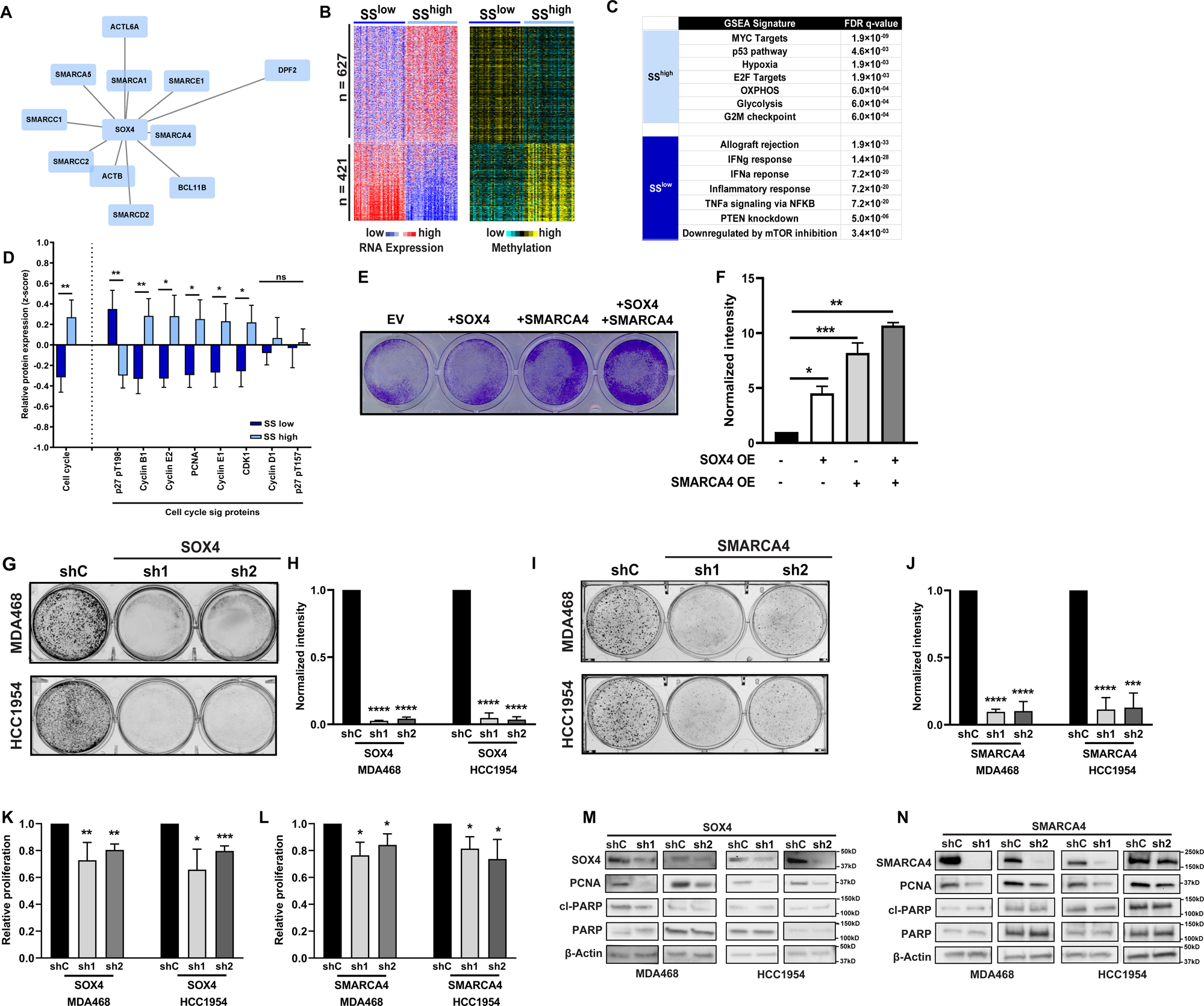
SOX4 and SMARCA4 regulate proliferation in basal-like breast cancer. (A) Plot of IP-MS data identifying interactions between SOX4 and SWI/SNF complex members; edge length is inversely proportional to the ratio of the spectra counts (B) Heatmaps depicting common differentially expressed (q<0.05) and methylated (q<0.01) genes in basal-like tumors with high (SS^high^) or low (SS^low^) SOX4 and SMARCA4 expression (C) GSEA analyses of differentially expressed genes (D) RPPA analysis of SS^high^ and SS^low^ basal-like tumor subpopulations identifies differentially expressed cell cycle proteins and a protein expression signature (E) SOX4 and/or SMARCA4 overexpression increases proliferation in MCF10A cells (F) quantification of crystal violet staining (G) ShRNA-mediated silencing of SOX4 (sh1, sh2) results in reduced colony formation in MDA-MB-468 or HCC1954 cells compared to scrambled control (shC) (H) Quantitation of colony formation (I) SiRNA-mediated silencing of SMARCA4 decreases colony formation in MDA-MB-468 and HCC1954 cells compared to control siC, quantified in (J). (K) Silencing of SOX4 and (L) SMARCA4 reduces MDA-MB-468 and HCC1954 cell proliferation (MTT assay) (M) shRNA-silencing of SOX4 reduces SOX4 and PCNA protein expression in MDA-MB-468 and HCC1954 cells; no effect was noted for total or cleaved PARP (N) Western blot analyses demonstrate decreased SMARCA4 and PCNA expression upon siRNA-mediated silencing of SMARCA4 in MDA-MB-468 and HCC1954 cells; no effect on total or cleaved PARP was observed (*p<0.05, **p<0.01, ***p<0.001 and ****p<0.0001.

To examine the potential role of the SOX4-SMARCA4 (SS) complex, we characterized basal-like tumors with high and low SOX4 and SMARCA4 gene expression. A Pearson correlation was first used to identify genes that were significantly and reproducibly correlated with SOX4 or SMARCA4 expression in basal-like tumors from the TCGA (n=185) and METABRIC (n=331) datasets (Figure S1A). These analyses identified a core set of 317 genes that were consistently positively (n=60) or negatively (n=257) associated with SS expression (Table S1). Consensus Cluster Plus [34] was then used to identify three distinct tumor subgroups defined by a high, moderate or low SS complex gene expression profile (Figure S1B-S1C). As illustrated for the TCGA (Figure S1D) and METABRIC datasets (Figure S1E), SS^high^ tumors were significantly enriched for the basal-like 1 (BL1) and mesenchymal (M) TNBC subtypes [39] while SS^low^ tumors were enriched for mesenchymal stem-like (MSL) and immunomodulatory (IM) subtypes (p<0.01), suggesting SS complex signaling is not characteristic of a given TNBC subtype.

To further characterize subgroups defined by SS complex activity, we examined differences in global gene and methylation patterns using orthogonal data from TCGA basal-like tumors (Figure S1F). Two-class SAM analyses [35] identified 628 (SS^high^) and 421(SS^low^) genes, respectively, that were activated at the mRNA level and exhibited decreased methylation in each subgroup (Figure 1B; Table S4). Gene Set Enrichment Analysis (GSEA) [36] was then used to assess pathway enrichment in each group. We determined that tumors with high SS complex expression were characterized by increased proliferation or cell cycle pathway signatures (i.e. Myc, E2F and G2/M) as well as metabolic pathways, including glycolysis and oxidative phosphorylation (OXPHOS) (Figure 1C). In contrast, tumors with low SS complex expression were defined by low mTOR or Akt signaling which is consistent with our previous studies reporting that SOX4 and SMARCA4 mediate PI3K/Akt signaling in basal-like breast cancer [15, 24]. To confirm the association between high SS complex expression and increased cell growth, protein expression was assessed in basal-like tumors (n=160) using TCGA Reverse Phase Protein Array (RPPA) data. SS^high^ tumors demonstrated increased expression (p<0.05) of proliferative proteins (i.e. Cyclin B1, Cyclin E2, PCNA, Cyclin E1 and CDK1) and a cell cycle protein signature (p=0.0095) [37] (Figure 1D), suggesting that SOX4 and SMARCA4 may contribute to aberrant tumor cell growth.

### SOX4 and SMARCA4 are required for basal-like cell growth

To examine the effect of SOX4 and SMARCA4 on cell growth, MCF10A cells were transduced to overexpress each protein alone or in combination. As shown in Figure 1E and 1F, when compared to an empty vector (EV) control, SOX4 overexpression increased proliferation 4.9-fold (p=0.0006), SMARCA4 overexpression increase proliferation 7.7-fold (p=0.0002) and overexpression of both proteins increased proliferation 10.7-fold (p<0.0001). To validate these data, we assessed the effect of shRNA-mediated silencing of SOX4 on cell growth and clonogenicity using basal-like HCC1954 and MDA-MB-468 cell lines which have high SOX4 and SMARCA4 mRNA and protein expression (Figure S2A-S2E). We determined that shRNA-mediated silencing of SOX4 using two unique shRNA (sh1 or sh2) resulted in a 54.5% - 65.5% reduction in SOX4 protein expression (Figure S3A) and a concomitant 95.5% - 97.5% reduction in colony formation (p<1.54×10^-08^) relative to the scrambled control in MDA-MB-468 cells (Figure 1G and 1H). Likewise, silencing of SOX4 in HCC1954 cells resulted in a 34.8% - 71.7% reduction in SOX4 protein levels (Figure S3B) which led to a 96.4% - 97.0% reduction in colony formation (p<1.6×10^-06^) (Figure 1G and 1H). Consistent with the proposed cooperative role of SMARCA4, we observed a 97.5% to 97.8% (p<3.4×10^-09^) decrease in colony formation following SMARCA4 silencing in MDA-MB-468 and HCC1954 cells, respectively (Figure 1I-1J, Figure S3C-S3D). As expected, shRNA-mediated silencing of either protein resulted in a ∼20-40% (p<0.01) reduction in short-term (96hr) cell growth in each cell line when assessed by MTT assay (Figures 1K-1L). Importantly, we observed a 39.5% - 79.7% (MDA-MB-468) and 18.3% - 75.6% (HCC1954) decrease in PCNA expression following shRNA-mediated silencing of SOX4 (Figure 1M) as well as a 47.2% (p=0.003) and 28.2% (p=0.007) decrease in PCNA expression following SMARCA4 silencing (Figure 1N) in each cell line. However, no change in PARP or cleaved PARP expression was observed (Figures 1M-1N, Figure S3A-S3D), suggesting that SOX4 and SMARCA4 are required to maintain cell growth and that reduced proliferation rather than apoptosis is responsible for decreased colony formation and cell growth.

### SOX4 and SMARCA4 function to cooperatively regulate glycolysis

Given the relationship between the SS complex and expression of metabolism genes in basal-like tumors, we next investigated the impact of SOX4 on cell metabolism. To do so, we overexpressed SOX4 in MCF10A cells (Figure S3E) and assessed changes in metabolite levels by liquid chromatography-mass spectrometry (LC-MS) (Table S5). SOX4 overexpression increased levels of glycolytic metabolites including glucose-6-phosphate (G6P, p=0.0007), glyceralehyde-3-phosphate (GADP; p=0.0002), dihydroxyacetone phosphate (DHAP; p=0.0008), 2-phosphoglycerate (2-PG; p=0.03), pyruvate (Pyr; p=0.006) and lactate (Lac; p=0.002) and decreased intracellular glucose (Glu; p=0.002) levels (Figure 2A-2B). Importantly, overexpression of SOX4 (p=0.003) or HA-tagged SOX4 (p=0.007) increased the glycolytic capacity of MCF10A cells while siRNA mediated silencing of SMARCA4 led to decreased (p=0.0003) ECAR (extracellular acidification rate) (Figure 2C).

**Figure 2.**
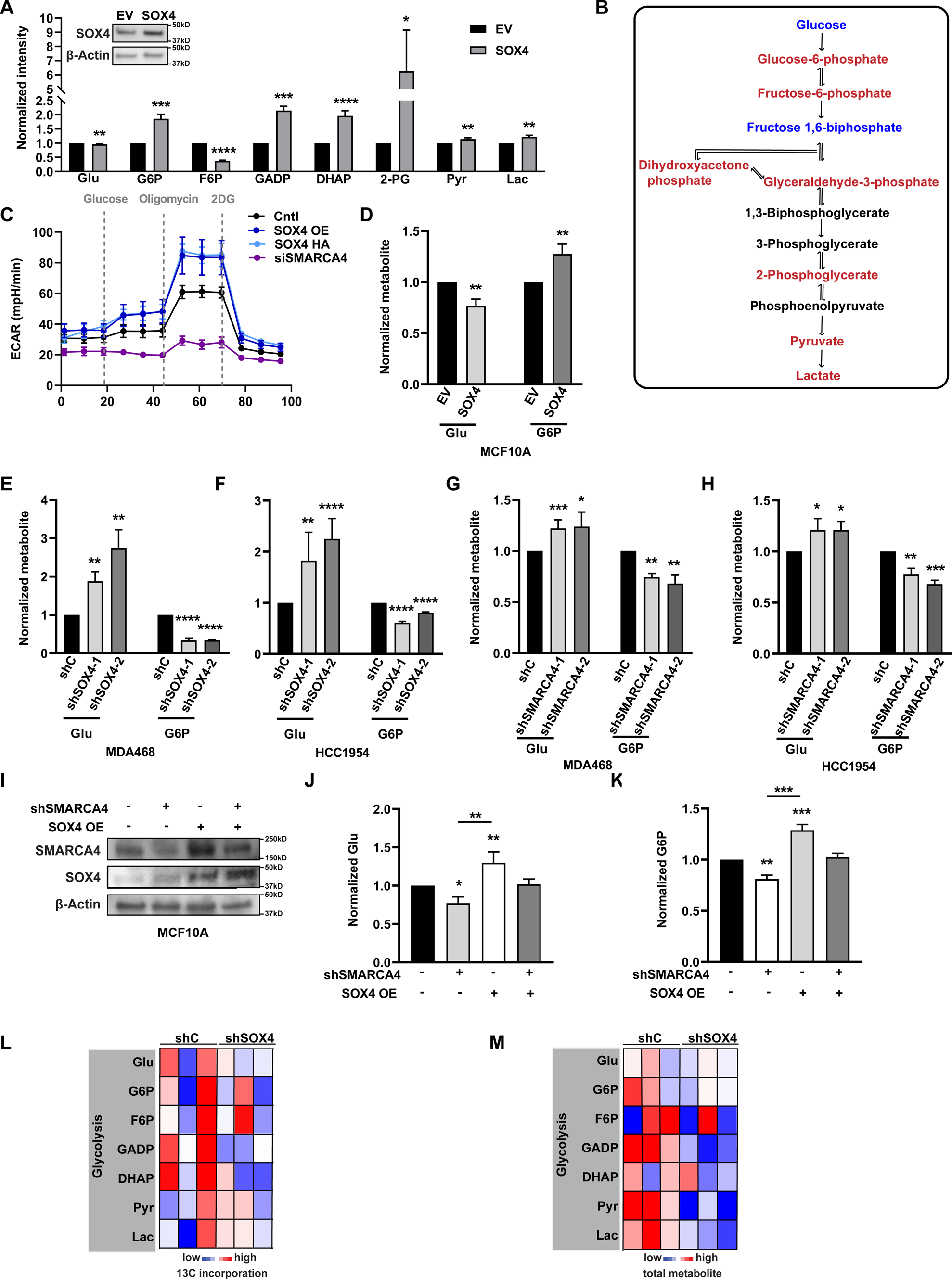
SOX4 and SMARCA4 mediate glycolysis in basal-like cell lines. (A) SOX4 overexpression increases glycolytic metabolites, as determined by LC-MS, compared to empty vector (EV) control transduced MCF10A cells; SOX4 Western blot (inset) is shown (B) Affected metabolites are mapped to the glycolysis pathway; up (red) or down (blue)-regulated metabolites are indicated (C) SOX4 or HA-SOX4 overexpression increases glycolytic capacity while silencing of SMARCA4 decreases ECAR in MCF10A cells (D) SOX4 overexpression decreases intracellular glucose and increases glucose-6-phosphate (G6P) in MCF10A cells (E) shRNA silencing of SOX4 increases intracellular glucose and decreases G6P in MDA-MB-468 cells or (F) HCC1954 cells compared to scrambled control (shC) (G) siRNA silencing of SMARCA4 increases intracellular glucose and decreases G6P in MDA-MB-468 or (H) HCC1954 cells compared to control (siC) (I) Western blot analyses demonstrate changes in SOX4 and SMARCA4 expression (J) Analysis of intracellular glucose and (K) G6P following SOX4 overexpression and/or silencing of SMARCA4 in MCF10A cells. (L) ^13^C glucose labeling of metabolites in MDA-MB-468 cells upon SOX4 knockdown demonstrates a decrease in glycolytic flux (M) Pooled metabolite analyses demonstrate decreased glycolytic pathway metabolites normalized to cell count *p<0.05, **p<0.01 and ***p<0.001.

We next examined the impact of SOX4 on glycolysis. SOX4 overexpression resulted in a 20.0% decrease in intracellular glucose (p=0.004) and a 1.2-fold increase in G6P levels (p=0.008) in MCF10A cells (Figure 2D). Consistent with these data, shRNA-mediated silencing of SOX4 increased intracellular glucose 1.8 – 2.1 (p<0.004) fold and reduced G6P expression 63.9% – 65.4% (p<1.0×10^-07^) in MDA-MB-468 cells (Figure 2E). HCC1954 cells showed a similar 1.8 – 2.3-fold increase (p<0.004) in intracellular glucose levels and a 19.9% – 40.0% decrease in G6P (p<1.6×10^-5^) following SOX4 silencing (Figure 2F).

Consistent with the proposed role of SMARCA4 as a SOX4 co-factor, siRNA-mediated silencing of SMARCA4 resulted in a 1.3-fold increase (p<0.05) in intracellular glucose levels and a 28.9% decrease (p=0.006) in G6P (Figure 2G) in MDA-MB-468 cells. A similar 1.2-fold increase in glucose (p=0.025) and a 46.3% decrease (p=1.3×10^-06^) in G6P (Figure 2H) was apparent in HCC1954 cells following SMARCA4 silencing. To confirm that SS proteins cooperatively mediate the glucose to G6P conversion, the impact of concurrent siRNA-mediated silencing of SMARCA4 and SOX4 overexpression was examined in MCF10A cells. As expected, SMARCA4 silencing resulted in a 1.3-fold increase (p=0.004) in intracellular glucose levels while SOX4 overexpression increased glucose consumption as evident by a 31.0% (p=0.0003) decrease in intracellular glucose (Figure 2I-2J). Importantly, concurrent SOX4 overexpression and siSMARCA4 silencing resulted in a phenotypic rescue of glucose consumption (p=0.06). Similar effects on G6P levels were observed in MCF10A cells as siRNA-mediated silencing of SMARCA4 led to a 40.2% reduction in G6P (p=0.008), SOX4 overexpression increased G6P 1.7-fold (p=0.03) while concurrent SOX4 overexpression and SMARCA4 silencing resulted in a phenotypic rescue of these cells (p=0.70) (Figure 2K). We do note that SOX4 overexpression leads to increased SMARCA4 protein levels in these experiments which is consistent with our recent observation that SOX4 can regulate the expression of its cofactor [24].

Finally, we performed in vitro glycolytic flux analysis using uniformly 13-carbon (^13^C) labeled [U^13^C6]-glucose to determine the impact of SOX4 silencing on MDA-MB-468 metabolic activity. Our analyses determined that SOX4 silencing resulted in decreased ^13^C labeling of glycolytic metabolites, including G6P (Figure 2L, Table S6). Consistent with our earlier analyses (Figure 2A), we observed a decrease in total glycolytic metabolite pools following SOX4 silencing (Figure 2M, Table S7).

### SOX4 and SMARCA4 modulate glycolysis through regulation of Hexokinase 2

Hexokinase (HK) proteins, of which HK2 is the most predominant in basal-like breast cancer, catalyze the glucose to G6P conversion [13, 47–50]. Given that we observed a shift in total G6P levels and ^13^C labeling in response to SOX4 or SMARCA4 manipulation (Figure 2) and that HK2 was identified as a component of the glycolysis pathway which showed increased expression in SS^high^ basal-like tumors (Figure 1), we speculated that changes in SS complex-dependent glycolytic activity may be due to altered HK2 expression. Consistent with this premise, we determined that SS complex expressing MDA-MB-468 and HCC1954 have high HK2 expression whereas MCF10A cells with lower SS expression have low HK2 levels (Figure S2F). As shown in Figure 3A, (quantified in Figure S4A), SOX4 overexpression increased HK2 protein levels 1.7-fold (p=0.03) in MCF10A cells. Likewise, shRNA-mediated silencing of SOX4 resulted in a 28.4% – 39.4% decrease in HK2 protein expression (p<0.05) in MDA-MB-468 cells (Figure 3B, Figure S4B) and a 50.3% – 86.8% decrease (p<0.002) in HCC1954 cells (Figure 3C, Figure S4C). SiRNA depletion of SMARCA4 resulted in a similar 65.3% (p=0.0003) and 44.5% (p<0.05) reduction in HK2 expression in MDA-MB-468 (Figure 3D, Figure S4D) and HCC1954 cells (Figure 3E, Figure S4E), respectively.

**Figure 3.**
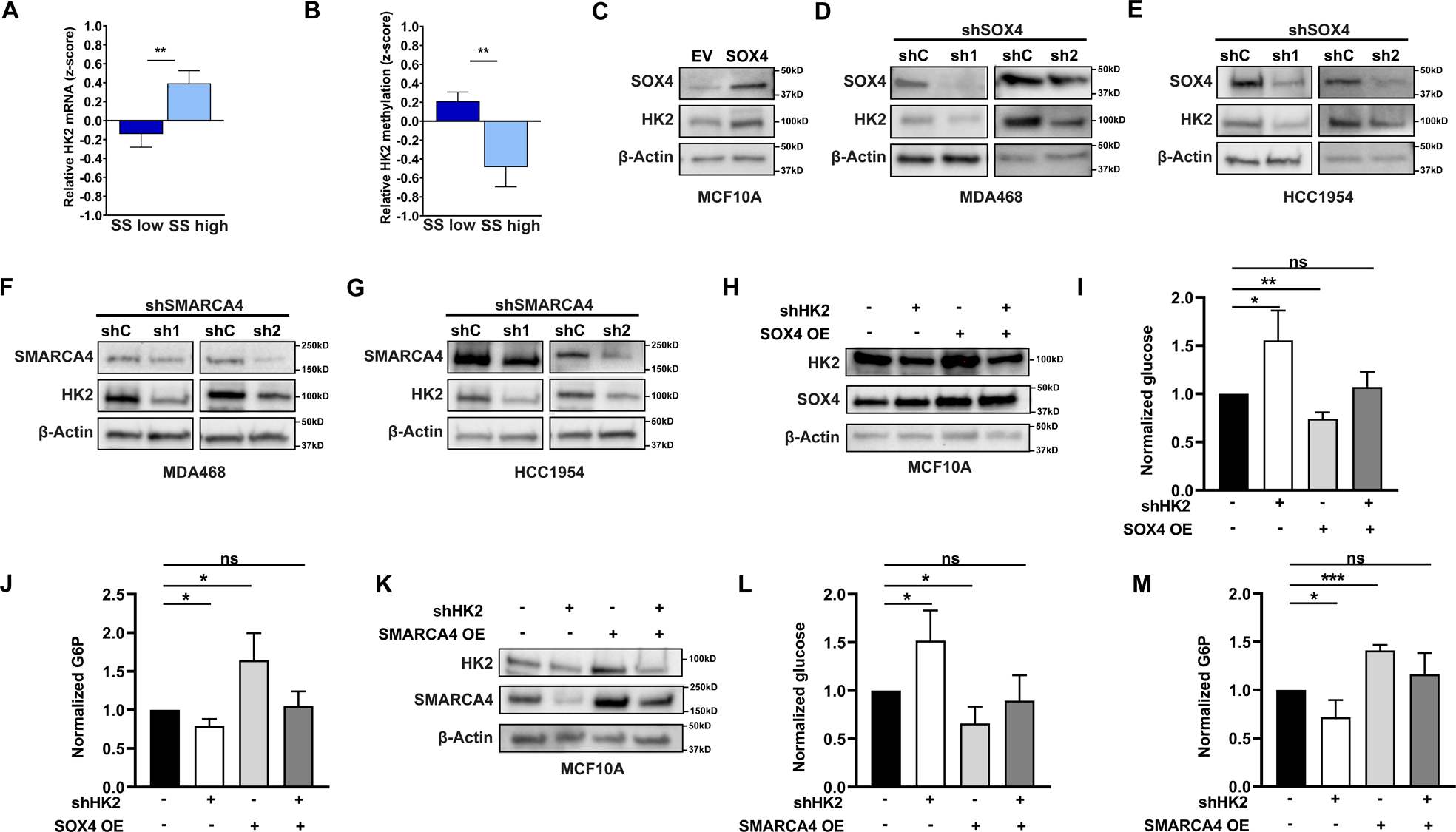
SOX4 and SMARCA4 modulate glycolysis through HK2. (A) Western blot analyses demonstrate SOX4 expression increases HK2 protein levels in MCF10A cells compared to empty vector (EV) control (B) shRNA silencing of SOX4 reduces HK2 protein expression in MDA-MB-468 or (C) HCC1954 cells compared to scrambled control (shC) (D) Western blot analyses demonstrate siRNA silencing of SMARCA4 decreases HK2 protein expression in MDA-MB-468 or (E) HCC1954 cells compared to scrambled control (siC) (F) Western blot analyses demonstrate changes in SOX4 and HK2 expression in MCF10A cells (G) Analysis of glucose and (H) G6P levels following SOX4 silencing and/or HK2 overexpression decreases upon HK2 knockdown and increases upon SOX4 overexpression (I) Western blot analyses demonstrate changes in SMARCA4 and HK2 expression in MCF10A cells (J) Analysis of glucose and (K) G6P levels following SMARCA4 overexpression and/or HK2 silencing *p<0.05, **p<0.01 and ***p<0.001.

To demonstrate that altered HK2 is responsible for aberrant SOX4-SMARCA4-mediated glycolysis, we assessed the impact of concurrent manipulation of HK2 and SOX4 or SMARCA4 on glucose and G6P levels. We determined that HK2 loss led to a 1.7-fold accumulation of intracellular glucose (p=0.04) while SOX4 overexpression reduced MCF10A glucose levels by 23.0% (p=0.002) (Figure 3F-3G). Importantly, concurrent SOX4 overexpression and HK2 silencing prevented SOX4-induced changes in glucose conversion (p=0.49). As expected, the inverse effect was observed for G6P levels, with HK2 silencing leading to a 23.4% decrease in G6P (p=0.02), SOX4 overexpression increasing G6P levels 1.7-fold (p=0.03), and concurrent SOX4 overexpression with HK2 silencing restoring G6P to basal levels (p=0.66) (Figure 3H).

Finally, to confirm that SMARCA4 is required to regulate this process, intracellular glucose and G6P levels were examined in MCF10A cells following manipulation of HK2 and/or SMARCA4. As expected, HK2 silencing increased intracellular glucose levels 1.5-fold (p=0.02), SMARCA4 overexpression led to a 44.0% decrease (p=0.03), while concurrent SMARCA4 overexpression with HK2 silencing restored intracellular glucose to basal levels (p=0.53) (Figure 3I-3J). Likewise, HK2 knockdown decreased G6P levels 28.7% (p<0.05), SMARCA4 overexpression increased G6P levels 1.4-fold (p=0.0002) and concurrent manipulation of HK2 and SMARCA4 restored (p=0.27) G6P to basal levels (Figure 3K). These data indicate that SOX4 and SMARCA4 promote the glucose to G6P conversion in an HK2-dependent manner.

### SOX4 and SMARCA4 directly regulate HK2 mRNA expression

Given that SOX4 and SMARCA4 are transcription co-factors, we examined the impact of this complex on HK2 mRNA expression [24]. We determined that shRNA-mediated silencing of SOX4 led to a 62.5% – 65.0% decrease in SOX4 mRNA expression (p<0.0002) and a concomitant 39.0% – 62.5% (p<0.003) decrease in HK2 mRNA levels in MDA-MB-468 cells (Figure 4A). This effect was confirmed in HCC1954 cells which demonstrated a 67.5% - 75% decrease in SOX4 mRNA expression (p<0.006) and a 48.8% – 79.0% decrease (p<0.006) in HK2 levels (Figure 4B). Importantly, we determined that SMARCA4 was required to mediate SOX4-dependent HK2 expression. As reported in Figure 4C, siRNA-mediated silencing of SMARCA4 resulted in a 48.0% decrease in HK2 mRNA levels (p=0.008), SOX4 overexpression led to a 2.1-fold increase (p=0.0001) while concurrent manipulation of SOX4 and SMARCA4 prevented SOX4-induced HK2 expression (p=0.08). These data indicate that the SOX4-SMARCA4 complex is required to cooperatively mediate HK2 transactivation.

**Figure 4.**
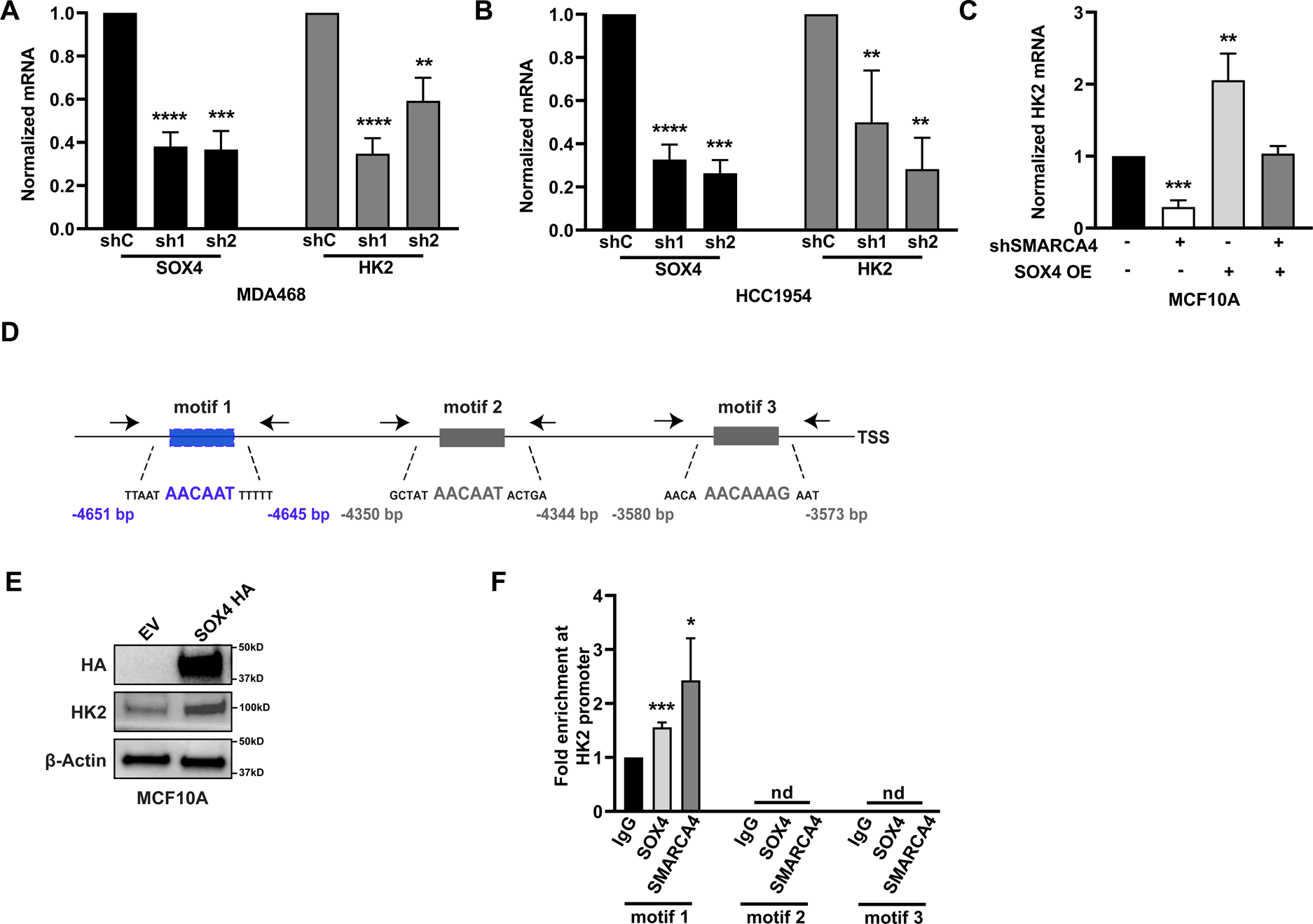
SOX4 and SMARCA4 regulate Hexokinase 2 expression by directly binding to its promoter. (A) shRNA-mediated silencing of SOX4 reduces HK2 mRNA expression (qRT-PCR) in MDA-MB-468 or (B) HCC1954 cells relative to scrambled control (shC) (C) SOX4 overexpression results in increased HK2 mRNA levels in a SMARCA4-dependent manner (D) Schematic of the HK2 promoter identifying three putative SOX4 binding motifs upstream of the HK2 transcription start site (TSS); forward and reverse primers are indicated by black arrows (E) Western blot analyses confirm increased HA-tagged SOX4 and HK2 expression in MCF10A cells following transduction of HA-SOX4 or empty vector (EV) control (F) ChIP-PCR demonstrates increased HA-SOX4 and SMARCA4 enrichment at the HK2 promoter compared to the IgG control at motif 1; no detected enrichment was observed at motif 2 or 3 (G) Western blot confirming SOX4 knockdown in HCC1954 cells (H) ChIP PCR demonstrates that in HCC1954 cells SOX4 is required for SMARCA4 enrichment on the HK2 promoter (I) SMARCA4 pulldown is comparable between the shC and shSOX4 transduced samples (J) Basal-like tumors (TCGA) with high SOX4 and SMARCA4 expression (SS^high^) show increased HK2 mRNA expression and (K) decreased methylation *p<0.05, **p<0.01, ***p<0.001 and ****p<0.0001.

Analyses of the regulatory region immediately 5’ of the HK2 Transcriptional Start Site (TSS) identified three regions which contained the canonical A/T-A/T-CAA-A/T-G SOX4 DNA binding motif [51] (Figure 4D). To identify the SOX4 binding motif, HA-tagged SOX4 was overexpressed in MCF10A cells, the increase in HK2 protein expression confirmed (Figure 4E) and ChIP qPCR analysis performed. Our analyses identified a significant 1.5-fold increase (p=0.0004) in HA-SOX4 enrichment at motif 1 (−4645bp) relative to IgG control (normalized to PNOC promoter as a negative control) while no detectable binding was observed at motif 2 (−4344bp) or motif 3 (−3573bp) (Figure 4F). Consistent with the role of SMARCA4 as a SOX4 co-factor, SMARCA4-specific ChIP-qPCR analyses identified a similar enrichment at motif 1 (p=0.03) suggesting that SOX4 and SMARCA4 cooperate to regulate HK2 mRNA expression (Figure 4F). Importantly, we demonstrate that SOX4 is required to recruit SMARCA4 binding at the HK2 promoter as shRNA-mediated silencing (Figure 4G) of SOX4 in HCC1954 cells reduced SMARCA4 enrichment by 73.3% when assessed by ChIP-qPCR (Figure 4H; p=0.0004). As evident by chromatin immunoprecipitation followed by SMARCA4 Western blot, SOX4 silencing does not reduce the ability of SMARCA4 to globally bind to chromatin (Figure 4I). These results are consistent with our previous work demonstrating that SOX4 is required to recruit SMARCA4 to the TGFBR2 promoter [24]. Consistent with these data, SS^high^ basal-like tumors demonstrate significantly higher HK2 mRNA expression (Figure 4J; q=0.007) and lower HK2 methylation (Figure 4K; q=0.003) suggesting that the described *in vitro* mechanism may be reflected in this subset of tumors.

### SOX4 and SMARCA4 regulate proliferation through Hexokinase 2

Given the noted role of glycolysis in promoting cell proliferation, we assessed whether SS complex driven cell growth is dependent on HK2 activity and glycolysis. We first determined that tumors with high SOX4 expression were characterized by increased SMARCA4 (p=8.1×10^-95^) and HK2 (p=2.6×10^-5^) mRNA levels. Likewise, these tumors showed increased proliferation (p=3.8×10^-24^), glycolysis (p=6.1×10^-11^) and glucose depletion (p=0.001) gene expression signatures (Figure 5A). Further analyses determined that average SOX4 and SMARCA4 expression was strongly associated (Spearman rank correlation) with proliferation (p=3.45×10^-73^) and glycolysis (p=1.25×10^-28^) signatures in human tumors; as expected a strong association was also evident between the proliferation and glycolysis (p=1.29×10^-118^) signatures. Basal-like tumors showed the highest levels of all three signatures (Figure 5B) which is consistent with the increased proliferation capacity of this subtype. Similar results were observed in the TCGA dataset (Figure S5A-S5B).

**Figure 5.**
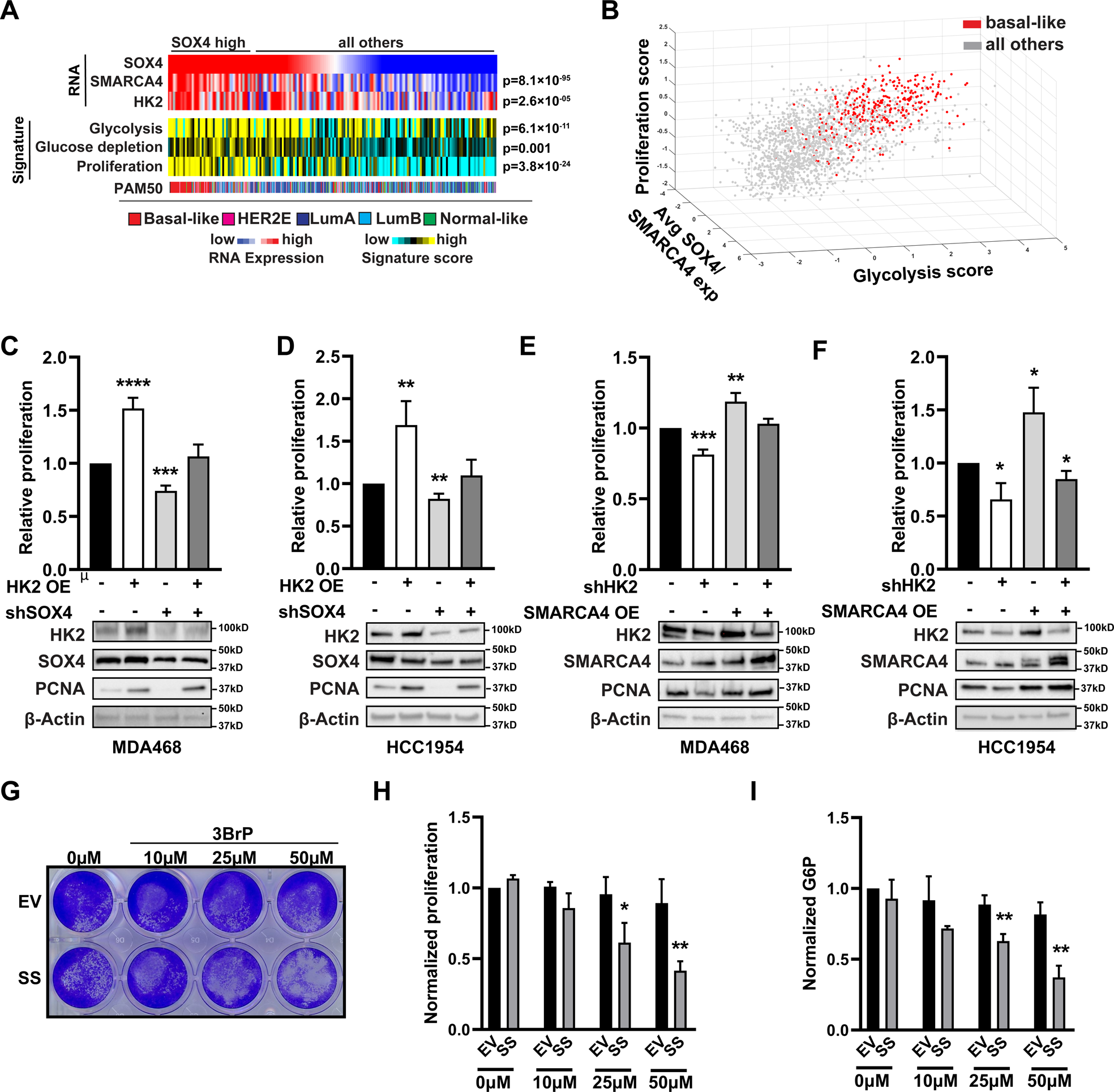
SOX4 and SMARCA4 regulate proliferation through Hexokinase 2. (A) The top quartile of SOX4 high samples from the TCGA dataset is enriched for SMARCA4 and HK2 expression and glycolysis-associated and proliferation signatures (B) Average SOX4 and SMARCA4 expression correlates with proliferation and glycolysis gene expression signatures in the METABRIC dataset; basal-like tumors are depicted in red (C) Relative proliferation (MTT) and protein expression (Western blot) demonstrate the effects of SOX4 and/or HK2 expression on MDA-MB-468 and (D) HCC1954 cell growth (E) Relative proliferation (MTT) and Western blots demonstrate the effects of SMARCA4 and/or HK2 expression on MDA-MB-468 and (F) HCC1954 cell growth (G) Proliferation assay of empty vector versus cells transduced with SOX4 and SMARCA4 lentivirus treated with different concentrations of 3-bromopyruvate, quantified in (H). (I) G6P measurements upon treatment with 3-bromopyruvate, confirming targeting of HK2 *p<0.05, **p<0.01, ***p<0.001 and ****p<0.0001.

Based on these data, we hypothesized that HK2 is necessary for SS complex-dependent proliferation. We determined that HK2 overexpression resulted in a 1.5-fold increase in proliferative capacity (p=4.6×10^-05^) of MDA-MB-468 (MTT assay) which was confirmed by a 1.2-fold increase (p=0.0005) in PCNA expression (Figure 5C, Figure S6A). Likewise, shRNA-mediated silencing of SOX4 (Figure S6B) in MDA-MB-468 cells resulted in a 23.0% (p=0.0009) decrease in cell proliferation and a 68.4% decrease in PCNA expression (p=0.02). Importantly, shRNA-mediated silencing of SOX4 with concurrent HK2 overexpression was able to rescue both phenotypes to wild-type levels (MTT, p=0.36; PCNA, p=0.72). Similar results were observed in HCC1954 cells (Figure 5D) where HK2 overexpression resulted in a 1.8-fold increase in cell growth (p=0.01) and 1.2-fold increase in PCNA expression (p=0.02) while shRNA-mediated silencing of SOX4 led to a 15.3% decrease in proliferation (p=0.007) and a 36.5% decrease in PCNA levels (p=0.02). Concurrent HK2 overexpression and shRNA-mediated silencing of SOX4 led to a phenotypic rescue (MTT, p=0.42) and no change in PCNA (p=0.35) relative to control cells (Figure S6C-S6D) indicating that HK2 is essential for SOX4 dependent cell growth.

We next examined whether SMARCA4-dependent basal-like cell growth is also dependent on HK2 activity. As shown, silencing of HK2 significantly reduced MDA-MB-468 cell proliferation (19.0%, p=0.0008) and PCNA expression (83.3%, p=0.005) while SMARCA4 overexpression promoted cell growth (1.2-fold increase, p=0.006) but did not significantly change PCNA levels (p=0.92) (Figure 5E, Figure S6E). As expected, concurrent silencing of HK2 with SMARCA4 overexpression restored the proliferative capacity (p=0.21) and PCNA expression (p=0.46) to basal levels. Similarly, HK2 silencing in HCC1954 cells resulted in a 36.1% (p=0.02) decrease in proliferation and a 44.1% (p=2.2×10^-05^) reduction in PCNA expression (Figure 5F, Figure S6F) while SMARCA4 overexpression increased proliferation 1.3-fold (MTT, p=0.02) and led to a modest, albeit not statistically significant, increase in PCNA (p=0.35). Similar to MDA-MB-468 cells, concurrent shRNA-mediated silencing of HK2 was able to prevent increased proliferation evident in SMARCA4 transduced cells as illustrated by an 18.0% reduction (p=0.03) in proliferation and no change in PCNA (p=0.96) relative to control cells.

Finally, we sought to demonstrate that SOX4-SMARCA4-dependent proliferation (Figure 1E-1F) is dependent on HK2 activity. To do so, MCF10A cells transduced with SOX4 and SMARCA4 (SS) or empty vector (EV) control (96h) were seeded at an equal density to a new plate and treated with increasing concentrations of 3-bromopyruvate (3BrP) for 24 hours (Figure 5G). Our analyses demonstrated SS transduced MCF10A showed a dose-dependent response to 3BrP treatment on cell growth (Figure 5H) and G6P levels (Figure 5I) with a 40.6% decrease in cell growth (p=0.01) and 49.2% decrease in G6P levels at the highest dose. Conversely, no effect on either phenotype was observed in EV transduced cells at any tested dose. Collectively, these data indicate that HK2 is required for SOX4 and SMARCA4-dependent proliferation.

## DISCUSSION

Triple negative breast cancer (TNBC) is an aggressive disease with few therapeutic options and poor clinical prognosis [3, 4, 30, 52]. Studies have indicated that altered glucose metabolism is a common feature of these tumors which contributes to TNBC development and/or progression [14, 49, 53–56]. Thus, identifying genetic mechanisms that promote metabolic reprogramming in these tumors will provide insight into TNBC or basal-like breast cancer genesis, progression and response to treatment.

SOX4 is one of the most commonly overexpressed genes in basal-like breast cancer and is associated with poor clinical outcome which underscores its potential oncogenic impact in these tumors [15–20, 23, 24]. Although SOX4 cannot efficiently transform cells or promote tumorigenesis alone, SOX4-dependent signaling is essential to maintain cell survival and to promote multiple oncogenic phenotypes [16, 17]. It was recently reported that SOX4 forms a complex with the SWI/SNF ATPase SMARCA4 which is essential for some aspects of SOX4 transcriptional activity, including regulation of PI3K/Akt signaling through altered TGFβR2 expression [24]. However, the global impact of the SS complex, which is highly expressed in basal-like breast tumors, on cellular signaling and tumorigenesis remains unclear.

In the current study, we demonstrate that differential expression of SOX4 and SMARCA4 is evident in subsets of basal-like tumors and that these subgroups are characterized by unique gene expression and methylation patterns that correspond with altered signaling networks. SS^high^ basal-like tumors demonstrate increased proliferation and metabolic reprogramming compared to SS^low^ basal-like tumors. Consistent with these data, multiple studies have reported that aggressive breast tumors are characterized by increased glycolysis compared to slower growing tumors or adjacent normal tissue [54, 56]. TNBC or basal-like cell lines demonstrate increased glycolytic capacity, increased proliferation and colony formation in high glucose medium and are sensitive to glycolytic inhibitors [12, 13, 49, 53–56] suggesting that increased glycolysis can stimulate TNBC cell growth and tumor progression. Despite these earlier studies, the mechanisms and signaling networks that promote metabolic reprogramming in these tumors remain unclear.

Using metabolomic analyses, we determined that SOX4 and SMARCA4 cooperatively promote glycolysis in basal-like cells lines in a HK2-dependent manner. Our studies demonstrate that the SS complex mediates this process through direct transcriptional activation of HK2. These results are consistent with tumor data showing a strong correlation between SOX4 and SMARCA4 expression, decreased HK2 methylation and increased HK2 expression in basal-like tumors. As such, our data support a model in which increased expression of the SS complex increases glycolytic capacity as a result of elevated HK2 expression and activity in a subset of basal-like tumors. We will note that SOX4 and SMARCA4 could mediate additional aspects of the glycolytic pathway and that other mechanisms may contribute to this process; however, these aspects of TNBC or basal-like tumor biology will require further study.

Given the noted link between increased glycolysis and the proliferative capacity of tumor cells [10, 11], we further determined that tumors with high SS complex expression show a strong association with increased proliferation and glycolysis. This was particularly evident in basal-like tumors which showed the highest levels of all three variables. Importantly, we have delineated a mechanistic link between SOX4-SMARCA4 mediated HK2 expression, glycolysis and cell growth. This is notable as previous studies have independently indicated that SOX4, SMARCA4 and HK2 are commonly expressed in basal-like or TNBC tumors and that each of these factors is essential for and/or can promote cell growth and survival [17, 25, 51, 57, 58]. As a result, our data define a novel signaling axis by which SOX4 and SMARCA4 cooperatively promote basal-like breast cancer cell growth through aberrant activation of HK2-dependent glycolysis. Additional studies have implicated SMARCA4 and HK2 in chemotherapeutic resistance and inhibition of immune activity suggesting that increased SS complex expression in basal-like breast tumors may have therapeutic implications [59, 60]. Importantly, our data suggest that the SOX4-SMARCA4 complex may regulate addition signaling in basal-like tumors; however, further work will be required to fully delineate this signaling network and its impact on this disease. Finally, SOX4 is commonly overexpressed in multiple forms of cancer, including prostate, melanoma and lung [22, 23, 61–67]. Given that SOX4 can regulate the expression of SMARCA4 [24], our data suggest that the SOX4-SMARCA4-HK2 signaling axis could represent a common oncogenic mechanism in other malignancies.

## Data availability

All data generated by genomic analyses and mass spectrometry has been included in the Supplementary Materials. Computational analyses were performed using publicly available data from the TCGA and METABRIC datasets as noted.

## Acknowledgements

We would like to thank members of our laboratory for their helpful suggestions and comments on this project.

