## Supplemental Figures for "SOX4-SMARCA4 complex promotes glycolysis-dependent TNBC cell growth through transcriptional regulation of Hexokinase 2"

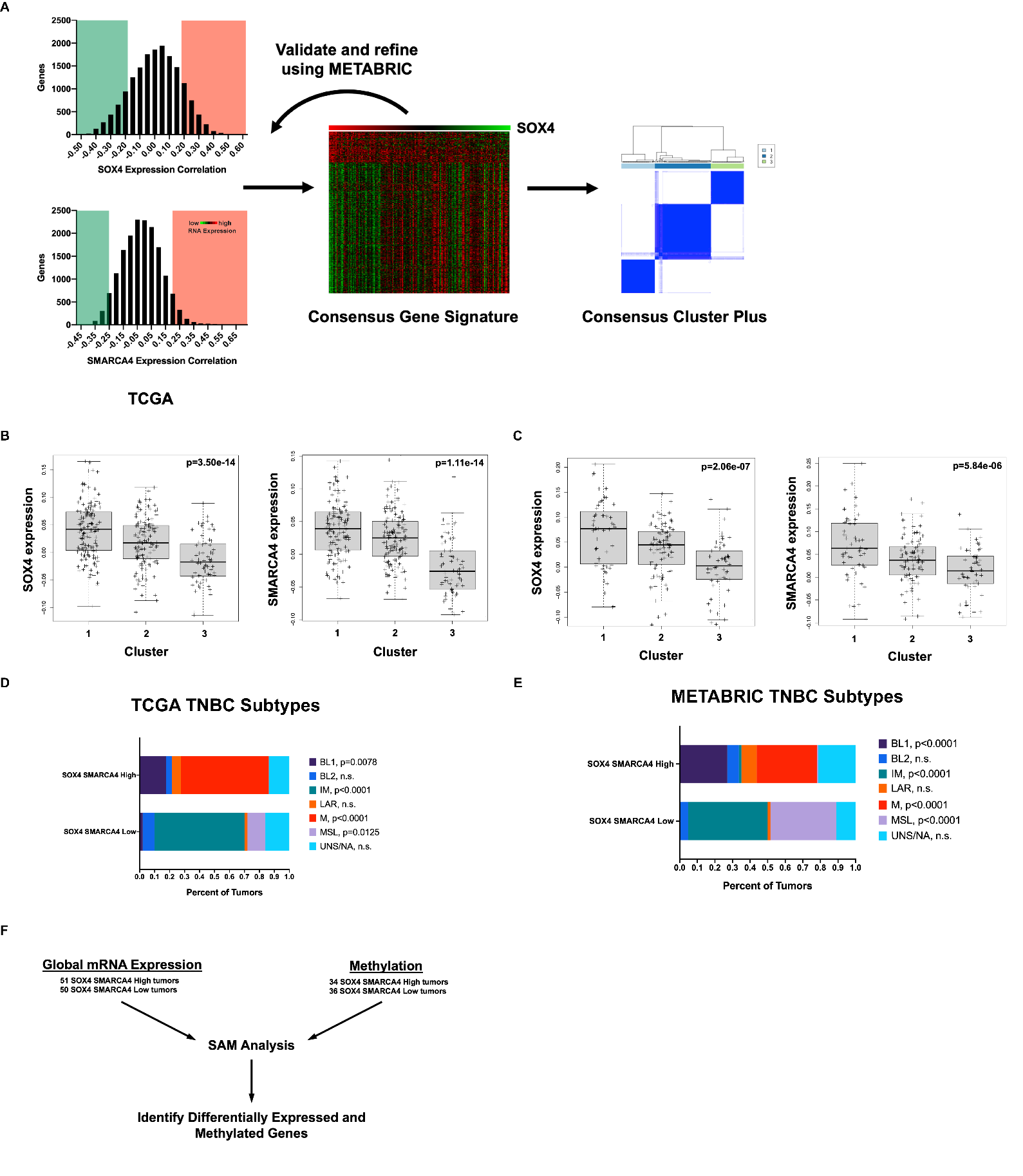

**Figure S1. Analysis of gene expression associated with high SOX4 and SMARCA4 expression.** (A) A Pearson correlation was used to identify genes that were consistently positive and negatively correlated (r>0.2, r<-0.2) with SOX4 and SMARCA4 using basal-like breast tumors (n=185) from the TCGA. These analyses were replicated in the METABRIC basal-like cohort (n=186) to generate a Consensus Gene Signature that corresponds with SOX4 and SMARCA4 expression. Consensus Cluster Plus was then used to define three clusters based on signature expression. Identified clusters are characterized by high (Cluster 1), moderate (Cluster 2) or low (Cluster 3) SOX4 and SMARCA4 expression in the (B) TCGA and (C) METABRIC datasets (D) Plot of TNBC subtypes from TCGA and (E) METABRIC datasets shows enrichment of the BL1 and M subtypes in the SOX4/SMARCA4 high samples and MSL and IM subtypes in the SOX4/SMARCA4 low subgroup (F) SAM analysis was used to assess gene expression and methylation data from SOX4/SMARCA4 high tumors (n=51) compared SOX4-SMARCA4 low tumors; a q<0.05 for gene expression and q<0.01 for methylation was used to define genes that showed increased mRNA expression (q<0.05) and decreased methylation (q<0.01) for each group. These analyses identified 628 and 421 genes that were characteristic of SS^high^ and SS^low^ complex basal-like tumors respectively.

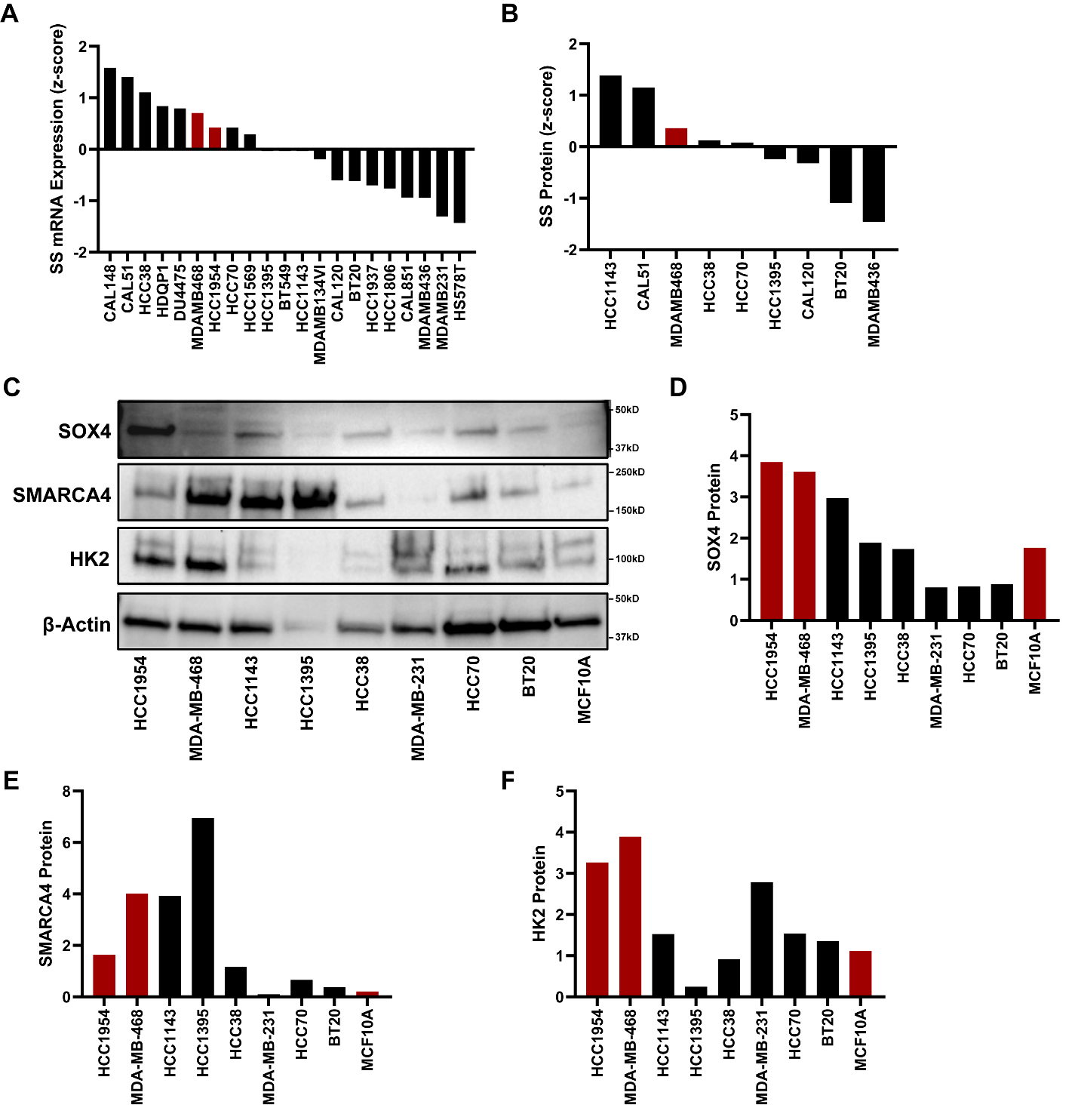

**Figure S2. Analyses of basal-like breast cancer cell lines identified MDA-MB-468 and HCC1954 cells with high SOX4, SMARCA4 and HK2 expression. (**A) Analysis of average (z-score) SOX4 and SMARCA4 mRNA (n=21) and (B) MS-derived protein expression (n=9) in basal-like breast cancer cell lines (n=21) from the Cancer Cell Line Encyclopedia (CCLE) dataset identified basal-like cells line with high or low SOX4 and SMARCA4 expression; protein data was not available for HCC1954 cells (C) Western blot analyses of a subset of basal-like cell lines confirms that MDA-MB-468 and HCC1954 cells have high SOX4, SMARCA4 and HK2 expression while MCF10A cells express lower levels of all three proteins. Western blot results are quantified for (D) SOX4 (E) SMARCA4 and (F) HK2.

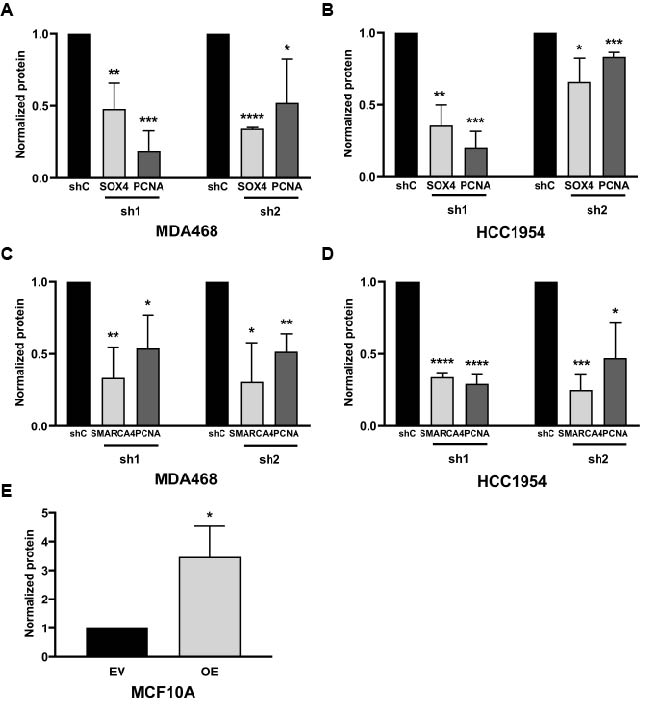

**Figure S3. Quantification of Western blots following SOX4 and SMARCA4 manipulation.** (A) SOX4 silencing results in 54.5% (sh1, p=0.007) and 65.5% (sh2, p=2.14×10^-08^) decrease in SOX4 protein expression in MDA-MB-468 cells and a 39.5% - 79.7% reduction in PCNA in MDA-MB-468 cells upon SOX4 knockdown (sh1, p=0.0006; sh2, p=0.05) compared to lentiviral control (shC) (B) shRNA-mediated silencing of SOX4 in HCC1954 cells results in a 71.7% and 34.8% decrease (sh1, p=0.001; sh2, p=0.021) in SOX4 protein levels and a 18.3% - 75.6% in PCNA (sh1, p=0.0003; sh2, p=0.0007) (C-D) SMARCA4 knockdown reduces SMARCA4 protein 72.1 – 82.8% in (C) MDA-MB-468 cells (p<0.01) or (D) 65.1 – 76.5% in HCC1954 cells (p<0.0002). PCNA levels decreased 37.0 – 42.7% in (C) MDA-MB-468 cells (p<0.03) and (D) 40.5 – 72.7% for HCC1954 cells (p<0.02) compared to control shRNA (shC) (E) Ectopic SOX4 expression increases SOX4 protein expression 2.7-fold (p=0.02) in MCF10A cells compared to empty vector controls (EV). * p<0.05, ** p<0.01 *** p<0.001 and **** p<0.0001.

­
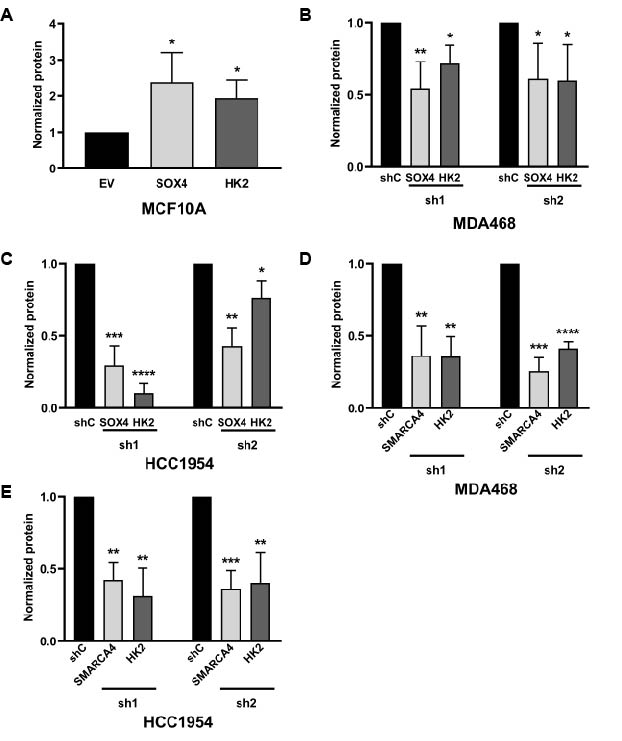

**Figure S4. Quantification of HK2 protein upon SOX4 and SMARCA4 manipulation.** (A) In MCF10A cells SOX4 overexpression increases HK2 protein levels compared to empty vector controls (EV). SOX4 silencing in (B) MDA-MB-468 and (C) HCC1954 cells using two different shRNA constructs resulted in reduced HK2 protein expression compared to non-silencing control shRNA transduced cells (shC) (D) SMARCA4 knockdown reduces HK2 protein compared to shRNA control (shC). For all studies: * p<0.05, ** p<0.01, *** p<0.001 and **** p<0.0001.

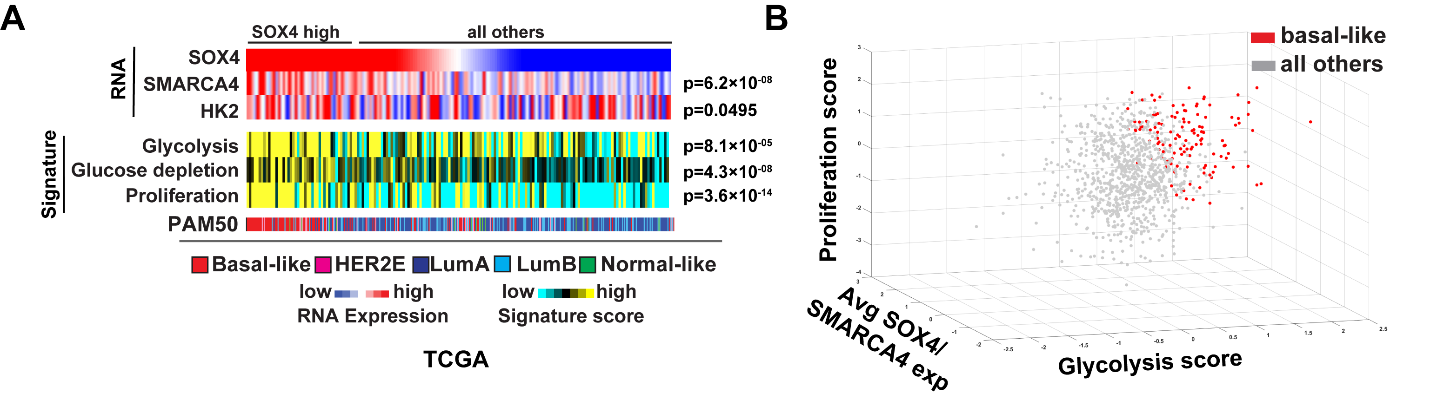

**Figure S5. SOX4 and SMARCA4 regulate glycolysis and proliferation in basal-like breast cancer samples from TCGA.** (A) SOX4 high samples (top quartile) are enriched for SMARCA4 and HK2 expression as well as glycolysis-associated and proliferation signatures in the TCGA dataset (B) The average of SOX4 and SMARCA4 expression correlates with both proliferation and glycolysis signatures in the TCGA dataset; basal-like tumors are highlighted in red and show a similar distribution relative to all three signatures.

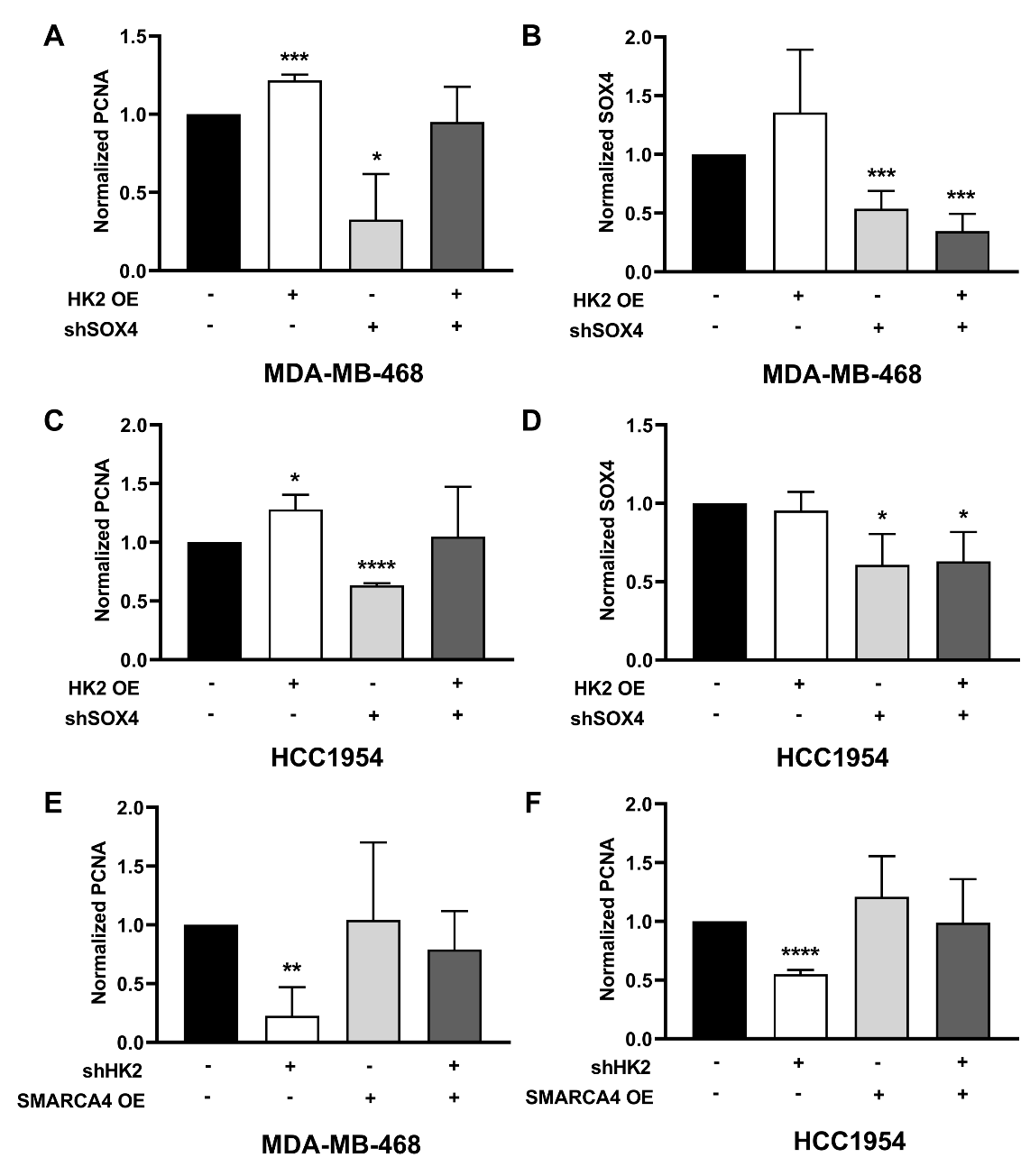

**Figure S6. Quantification of proliferation upon manipulation of SOX4, SMARCA4 and HK2.** (A) Quantification of Western blots for MDA-MB-468 cells shows an increase in PCNA expression upon HK2 overexpression, a reduction upon SOX4 knockdown and a return to levels comparable to control upon both (B) SOX4 quantification in MDA-MB-468 cells shows a reduction in protein expression upon SOX4 knockdown (C) Western blot of HCC1954 cells shows a similar pattern of PCNA and (D) SOX4 expression (E) MDA-MB-468 cells show a reduction in PCNA upon HK2 knockdown but no change upon SMARCA4 overexpression (F) Similar results were found in HCC1954 cells. * p<0.05, ** p<0.01, *** p<0.001, and **** p<0.0001.

**SUPPLEMENTAL TABLES**

**Table S1.** SOX4-SMARCA4 Consensus Gene Signature

**Table S2.** Summary of TCGA basal-like breast cancer samples

**Table S3.** Summary of METABRIC basal-like samples

**Table S4.** Summary of genes in the SS^high^ and SS^low^ basal-like tumor groups

**Table S5.** Summary of SOX4-induced metabolites in MCF10A cells

**Table S6.** Summary of ^13^C glucose labeled glycolysis metabolites upon SOX4 knockdown in MDA-MB-468 cells

**Table S7.** Summary of glycolysis metabolite values following SOX4 knockdown in MDA-MB-468 cells
